# Application of the Bradley-Terry model to quantify components of sperm competition

**DOI:** 10.64898/2026.07.06.736847

**Authors:** Mehrnaz Afkhami, Mona L. Li, Crystal Liang, Prajal H. Patel, Norene A. Buehner, Mariana F. Wolfner, Andrew. G. Clark

## Abstract

In many species with female sperm storage, ejaculates from multiple males overlap in the female reproductive tract, making sperm competitive ability a key component of male reproductive fitness and a target of rapid evolutionary change in the underlying genes. Here, we used controlled laboratory assays of *Drosophila melanogaster* sperm competition, with doubly-mated females and paternity assignment of offspring, to ask whether a Bradley-Terry framework can effectively summarize and predict competitive outcomes. The Bradley-Terry model is a probabilistic approach that estimates a latent “ability” score for each contestant based on outcomes of pairwise contests, and thus is naturally suited to data from sperm competition, which are intrinsically pairwise. We selected five distinct male genotypes: four carried strongly expressed RFP or GFP markers that allowed us to distinguish their heterozygous offspring under UV illumination, and the fifth was Canton-S, a standard wild-type genotype that served as our reference. Using Canton-S females, we assayed all 20 ordered pairwise combinations of first and second male, recorded successful double matings, and quantified the offspring sired by each male. We then extended the Bradley–Terry model to estimate genotype-specific competitive success separately for first-male “defense” (fertilization success following initial mating, also called “P1”) and second-male “offense” (fertilization success following a remating, also called “P2”). This framework provides a flexible and efficient way to integrate results across large arrays of pairwise mating tests and to derive predictive scores for sperm competitive performance.

## Introduction

Sperm competition, the competition between ejaculates of different males for fertilization within a multiply mated female, is now recognized as a pervasive and powerful component of sexual selection and male fitness across a broad range of taxa. In *Drosophila melanogaster*, experimental designs in which single females are mated sequentially to two males carrying distinguishable genetic markers have provided a central framework for quantifying “second male precedence” and related measures of competitive fertilization success. Yet even in this model system it has proved surprisingly difficult to obtain clean estimates of the underlying displacement processes. For example, Gilchrist and Partridge (2000) showed that variation in copulation duration strongly affects sperm transfer, complicating attempts to model sperm displacement from simple behavioral or paternity data alone and highlighting the sensitivity of standard assays to uncontrolled sources of variance.

More broadly, a retrospective synthesis of the “scientific revolution” in sperm competition research over the past fifty years emphasizes both the conceptual importance of sperm competition as a major, and often dominant, component of male fitness and the methodological challenges that have hindered precise, comparable measurement of its strength across studies and systems (Simmons and Wedell 2020). Given this dual reality, biological importance coupled with statistical and experimental difficulty, there is a clear need for quantitative frameworks that can leverage multiple pairwise mating tests to estimate the strength of sperm competition with greater rigor, and that offer genuine predictive accuracy for competitive outcomes; such approaches would substantially advance our ability to integrate postcopulatory processes into evolutionary and quantitative genetic models of fitness.

Sperm competition in *Drosophila melanogaster* has emerged as a powerful system for dissecting the genetic basis of postcopulatory sexual selection. Clark and colleagues showed extensive standing variation among male genotypes in their ability to compete against rival sperm, implicating segregating genetic factors that influence competitive fertilization success (Clark et al. 1995). Subsequent association studies identified polymorphism in accessory gland and other seminal fluid protein genes as key contributors to this variation, with allelic differences on both the second and third chromosomes affecting sperm competitive ability (Fiumera et al. 2005; Fiumera et al. 2007). These findings are consistent with broader evidence that variation in seminal fluid gene expression and function has substantial consequences for sperm competition outcomes and paternity success (Patlar and Civetta 2022; Patlar et al. 2023).

It is important to note that competitive outcomes are not always strictly hierarchical: male genotypes can exhibit nontransitive relationships, such that no single genotype consistently dominates all others, complicating the notion of a simple fitness ranking (Clark, Dermitzakis, and Civetta 2000). Similar complexity is evident in variation in sperm competition and sperm competition avoidance in natural populations (Civetta et al. 2008). These empirical results, together with comparative work on sperm heteromorphism and spermicide in Drosophila and other insects (Holman et al. 2008; Holman and Snook 2008), reinforce the view that postcopulatory sexual selection operates on a suite of male and female traits. Motivated by such data, theoretical analyses have shown that sperm competition can maintain genetic polymorphism, with allele frequency dynamics shaped by frequency dependence, genotype interactions, and mating structure, leading to complex and sometimes counterintuitive evolutionary trajectories (Clark 2002; Snook 2002).

In this paper, we develop and apply a Bradley-Terry framework for estimating sperm competitive abilities from pairwise mating assays. The Bradley-Terry model (Bradley & Terry, 1952) was originally devised to infer latent “abilities” of competing entities, such as sports teams, using only win-loss outcomes from pairwise contests, providing a natural way to recover a hierarchy of competitive quality even when the schedule of contests is incomplete and not all teams play one another. By mapping sperm competition outcomes in doubly-mated females onto this framework, we can treat each male genotype as a “team” and each mating trial as a contest, using the model to estimate continuous measures of sperm competitive ability and to generate probabilistic predictions for outcomes of male–male combinations that have not been directly tested. Building on this analogy, we employ both the standard Bradley-Terry formulation and an extension that partitions competitive ability into components corresponding to first-male (P1) and second-male (P2) precedence, allowing more refined inference about distinct aspects of sperm displacement and defensive performance.

## Materials and methods

### Stocks and Fly cultures

We maintained all Drosophila stocks on a standard yeast–glucose medium at 25° C under a 12 h light : 12 h dark cycle. We selected four lines with clearly distinguishable GFP or RFP expression patterns that were visible in adult heterozygotes, and included Canton-S as a fifth, wild-type line. Canton-S was designated as C. One GFP line, expressing in the eye and ocelli (23507, *w*^*1118*^; Mi{GFP[E.3xP3]=ET1}MB03040) (designated as E), was obtained from the Bloomington Drosophila Stock Center (https://bdsc.indiana.edu/), while the remaining lines showed whole-body GFP {Sqh-tdGFP}2R (designated as B), whole-body RFP {Sqh-mCherry}IV-1 (designated as A), or thorax-and-abdomen GFP expression (ubi-GFP;ProtamineB-GFP) (designated as U).

All experimental females were Canton-S, and Canton-S males served as the reference genotype in sperm competition assays. Each line was uniquely identifiable by the combination of marker color, expression pattern, and fluorescence intensity, and flies from all but one line had wild-type red eyes.

### Sperm competition assay

We implemented a 5 × 5 grid design (Figure 1) in which all pairwise combinations of male lines were assayed. There are 10 ways to draw pairs from these 5 lines, and each pair of male genotypes can be presented to females in two orders, giving a total of 20 experimental double matings. Virgin males and females from Canton-S, together with virgin males from the four fluorescently labeled lines, were collected and aged for 3–5 days prior to mating. For the first mating, a single male and female were placed together in a vial containing standard medium for 24 h, after which the male was removed. On the second day, the female was transferred to a fresh vial with a new male for an additional 24 h, then the second male was removed and the female was moved to a third vial on day 4. Thereafter, the female was transferred to new vials every other day for 8 days following the initial mating. Offspring were scored 12–14 days after egg laying under a dissecting microscope equipped with NightSea fluorescence illumination, and the number of progeny sired by each male was quantified from their fluorescent marker phenotypes. We set up 818 crosses and 330 were successfully doubly mated and sired more than 20 offspring. Replicate counts for each pairwise test were: (AB=11; AC=20; AE=16; AU=14; BA=12; BC=19; BE=18; BU=6; CA=14; CB=25; CE=20; CU=21; EA=27; EB=26; EC=19; EU=16; UA=8; UB=9; UC=13; UE=16)

**Figure 1.**
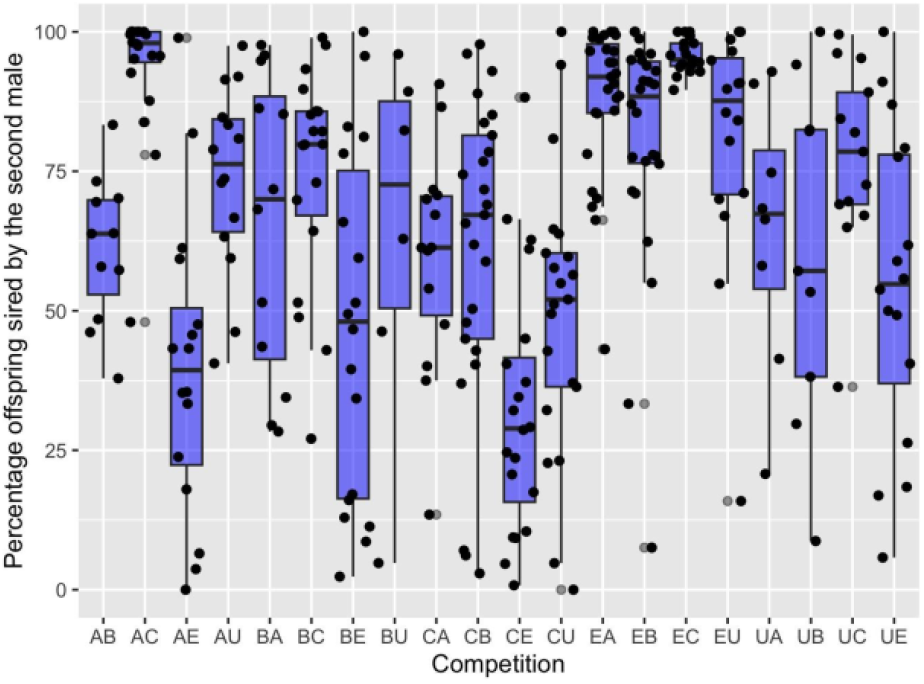
Percentage of offspring sired by the second male (P2) in each of the 20 crosses.

### Statistical analysis

All analysis was performed in R (version 4.5.1). The R packages employed included BradleyTerry2 (Turner & Firth, 2012), ggplot2 (Wickham, 2016), and tidyverse (Wickham et al., 2019).

### Modeling

In the original Bradley-Terry formulation, competing teams are not assumed to have any intrinsic order; the goal is simply to estimate an “ability” parameter for each team based on the outcomes of pairwise contests. Let the abilities of teams 1 and 2 be *A*_*1*_ and *A*_*2*_; under the model, the probability that team 1 wins a contest against team 2 is *A*_*1*_/(*A*_*1*_ + *A*_*2*_). With this assumption, Bradley and Terry (1952) derived maximum likelihood estimators for the ability parameters using the observed pattern of wins and losses across a series of contests. If one samples from a population in which many matings have occurred in all possible orders, and the order of mating is ignored, this same framework could, in principle, be applied to sperm competition data to yield a single net sperm competitive “ability” for each male line. However, such an approach would obscure a key biological feature of Drosophila sperm competition: mating order has a substantial effect on paternity, with roughly 80% of offspring typically sired by the second male, on average (Manier et al., 2013). As a result, an order-free Bradley-Terry model fails to capture an important axis of variation in competitive fertilization success.

### An order-dependent Bradley–Terry model

To accommodate the biological reality that mating order matters, we extend the Bradley-Terry framework to estimate two distinct sperm competition abilities for each line: one when the male is the first to mate (the “defense” or P1 component) and one when the male is the second to mate (the “offense” or P2 component). The R code TwoOrderBradleyTerry.R fits a binomial model to pairwise mating data, estimating two latent parameters for each male type: a **defensive** term *D*_*i*_ and an **offensive** term *O*_*i*_. For each double mating, it models the number of offspring sired by the first male, *y*_1_, out of the total *N* = *y*_1_ + *y*_2_, as *y*_1_ ∼ Binomial(*N, p*), with *p* = *D*_*P*1_/(*D*_*P*1_ + *O*_*P*2_), so the probability that the first male sires a given offspring increases with his defensive ability and decreases with the opponent’s offensive ability. The code parameterizes all effects on the log scale for optimization stability and identifiability. One male type is fixed as the reference by setting *D*_1_ = 1, while the remaining defensive effects and all offensive effects are represented as exponentiated free parameters, so they are constrained to be positive. The function *optim* minimizes the negative log-likelihood using the BGFS algorithm, and the fitted values are recovered by exponentiating the estimated log-scale parameters. Standard errors are estimated as the square root of the diagonal elements of the inverse Hessian matrix, so these are approximate asymptotic standard errors based on the local curvature of the likelihood surface at the optimum.

### Goodness-of -fit

The code also computes a goodness-of-fit deviance as 2(*l*_sat_ − *l*_model_), where *l*_sat_is the log-likelihood of a saturated model that fits each row exactly and *l*_model_is the fitted model’s log-likelihood. Smaller deviance means the fitted model is closer to the saturated, perfectly flexible model; larger deviance means worse fit. It then compares that deviance to a chi-square distribution with df = #assays - #parameters to get a *P*-value, which is intended as a rough test of whether the model fits substantially worse than a saturated model.

## Results

### Number of offspring

Because sperm competitive ability is inferred from offspring counts, the precision of these estimates depends in part on the total number of offspring produced. In these experiments, the four fluorescently labeled genotypes (A, B, U, and E) sired similar numbers of offspring in the first vial, whereas Canton-S produced more offspring than any of the others (Figure S1). Total offspring number also varied across competition assays (Figure S2).

### General sperm competition ability using the Bradley-Terry model

The 5×5 grid cross encompassed all 10 pairwise combinations of the five male genotypes, with each pairing performed in both mating orders (i.e., which male mated first versus second). For each cross, data consisted of the number of offspring sired by each male, with each female’s progeny constituting an independent replicate of the pairwise comparison (Figure 1). Note the large differences between the P2 values (proportion of offspring sired by the second male) of some of the reciprocal orderings, including AE vs. EA and CE vs. EC.

Fitting the standard Bradley–Terry model requires ignoring mating order, effectively treating both orders as equally frequent. The pooled data therefore capture the average offspring sired by each male genotype across mating sequences. The resulting Bradley–Terry estimates provide a net measure of sperm competitive ability for each genotype, normalized to Canton-S (Table 1).

**Table 1.** Estimates of the sperm competitive abilities under the model assuming that males engage in matings as the first vs. the second male equally often. Line A was considered the reference.

| Genotype | Ability | S.E. |
| --- | --- | --- |
| **C** | 0.368 | 0.018 |
| **B** | 0.018 | 0.019 |
| **A** | 0 | - |
| **U** | -0.028 | 0.020 |
| **E** | -0.981 | 0.019 |

Model fit appeared adequate, as the Q–Q plot showed no notable deviations from expectation (Figure S2). We next fit a model incorporating a “home advantage,” treating the second male as advantaged, consistent with double-mating assays in *D. melanogaster* in which the second male typically sires ∼80% of offspring (with substantial variation among lines). Incorporating this effect altered the relative ranking of the A and B genotypes (Table 2). The home-advantage model provided a significantly better fit than the base model (BT base AIC score = 21760 vs. BT with home advantage AIC score = 14407, so ΔAIC = 7353; *P* < 2.2 × 10^−16^;).

**Table 2.**
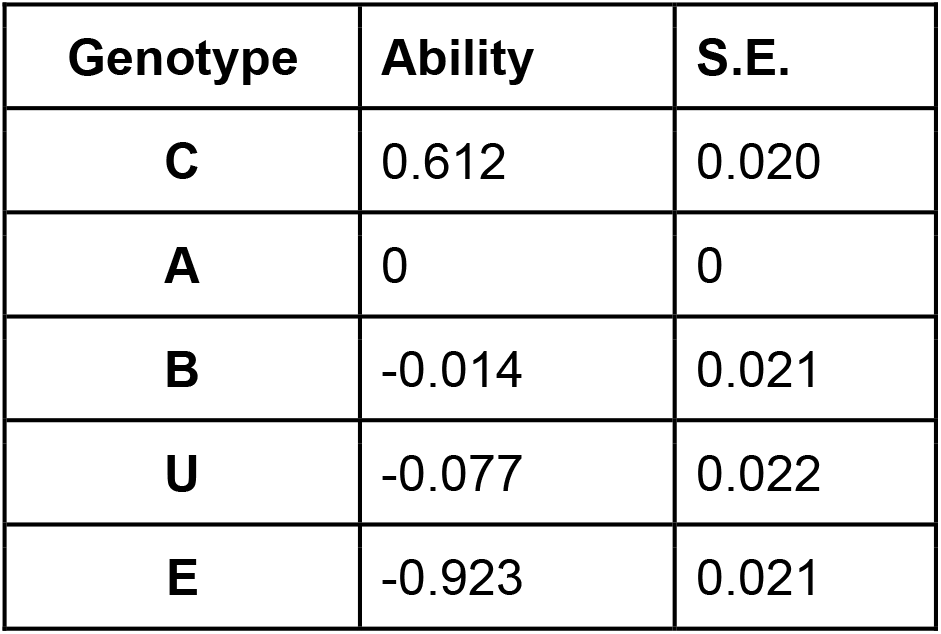
Sperm competitive ability estimates as in Table 1, but allowing for an additional “home advantage” parameter.

| Genotype | Ability | S.E. |
| --- | --- | --- |
| **C** | 0.612 | 0.020 |
| **A** | 0 | 0 |
| **B** | -0.014 | 0.021 |
| **U** | -0.077 | 0.022 |
| **E** | -0.923 | 0.021 |

**Table 3.** Defense and offense components of sperm competitive abilities estimated with the two-factor Bradley-Terry model described in this paper.

| Genotype | Defense (P1) | S.E. defense | Offense (P2) | S.E. offense |
| --- | --- | --- | --- | --- |
| **A** | 0 | — | 3.017 | 0.037 |
| **B** | 1.343 | 0.036 | 2.362 | 0.032 |
| **C** | 1.823 | 0.031 | 6.796 | 0.037 |
| **E** | 0.503 | 0.038 | 1.038 | 0.032 |
| **U** | 1.253 | 0.039 | 2.153 | 0.031 |

### The two-factor Bradley-Terry model

Because the experimental design recorded mating order, it is in principle possible to estimate both defensive (first-male) and offensive (second-male) components of sperm competition. Extending the Bradley–Terry model to distinguish between the two mating orders substantially improved model fit (Figure 2).

**Figure 2.**
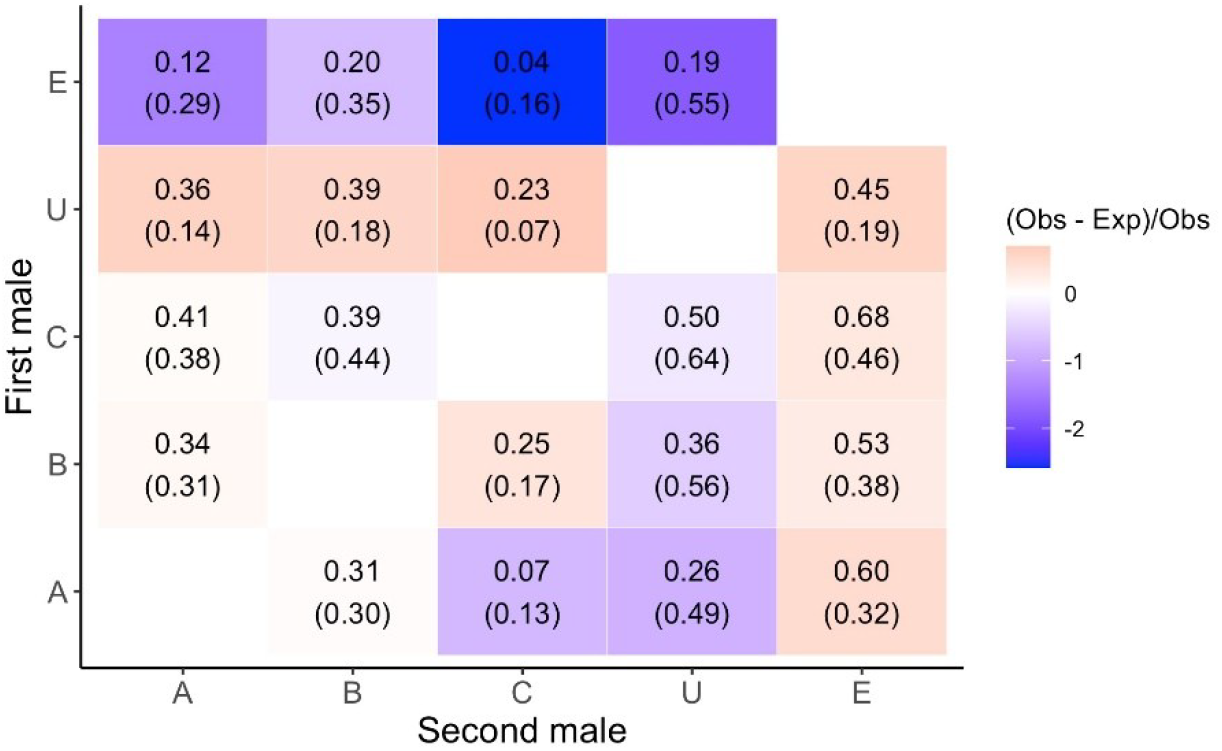
The observed and (expected) P1 defense abilities shown in the 5x5 pairwise grid, with the shading indicated scaled difference between the two.

This result is unsurprising given the strong effect of mating order on relative paternity success. The model yields separate estimates of offense and defense parameters for each line. Similar estimates have been obtained in studies using competition against a single marked reference line, and it has been observed that the offense and defense estimates are often positively correlated, as seen here (Figure 3).

**Figure 3.**
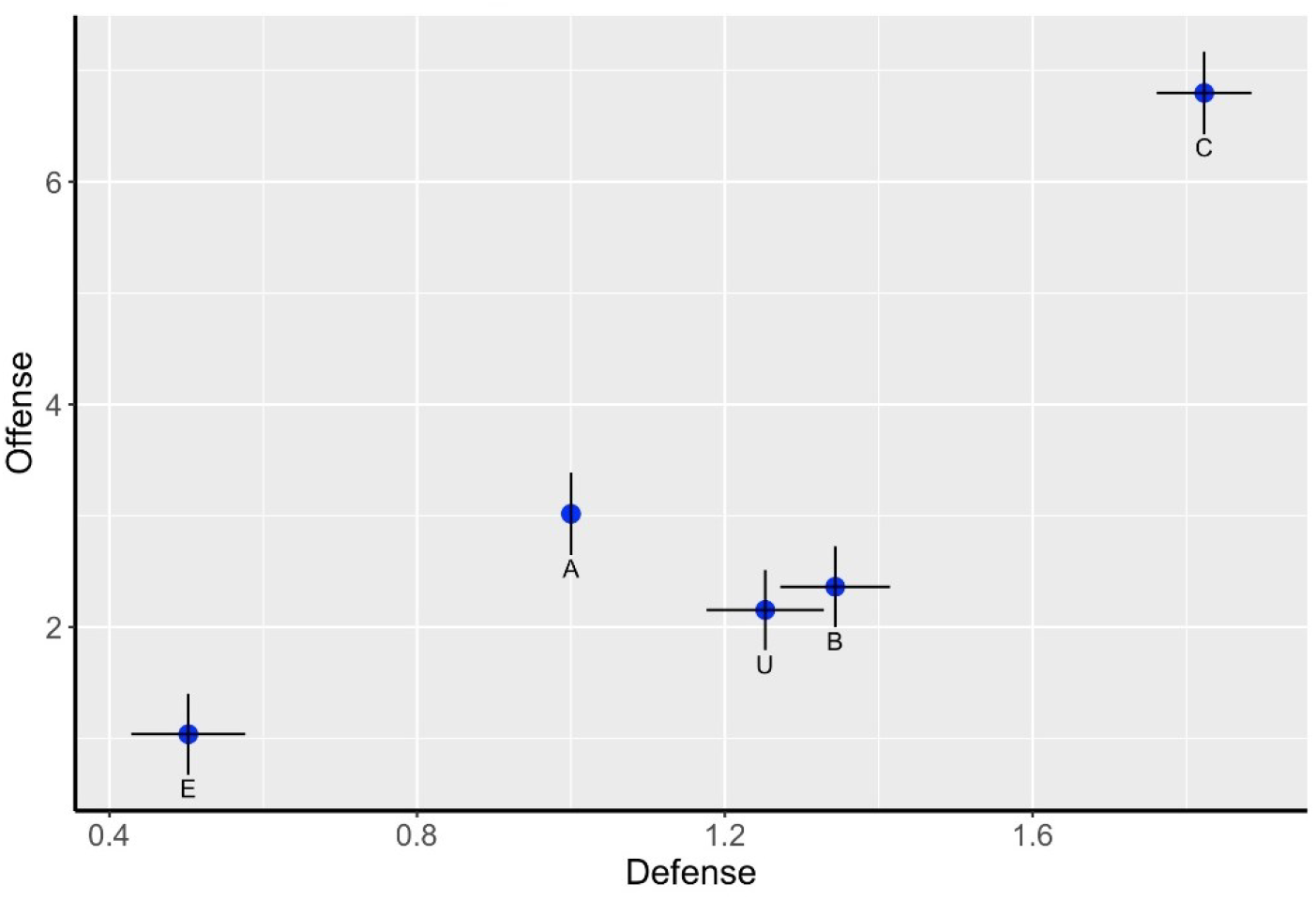
Estimated abilities for defense (P1) and offense (P2) along with standard errors. The five lines are clearly distinct in both P1 and P2, with lines U and B being similar in both sperm competition metrics.

To assess whether replication was sufficient to yield stable estimates of sperm competitive parameters, we performed a simple computational subsampling analysis. Random subsets of mating trials were drawn and the model refit to each subset. When sample sizes were small, the inferred estimates varied substantially, including changes in rank order; however, as sample size increased, the estimates stabilized. A practical criterion for adequate replication is stabilization of rank order, which is observed here (Figure 4).

**Figure 4.**
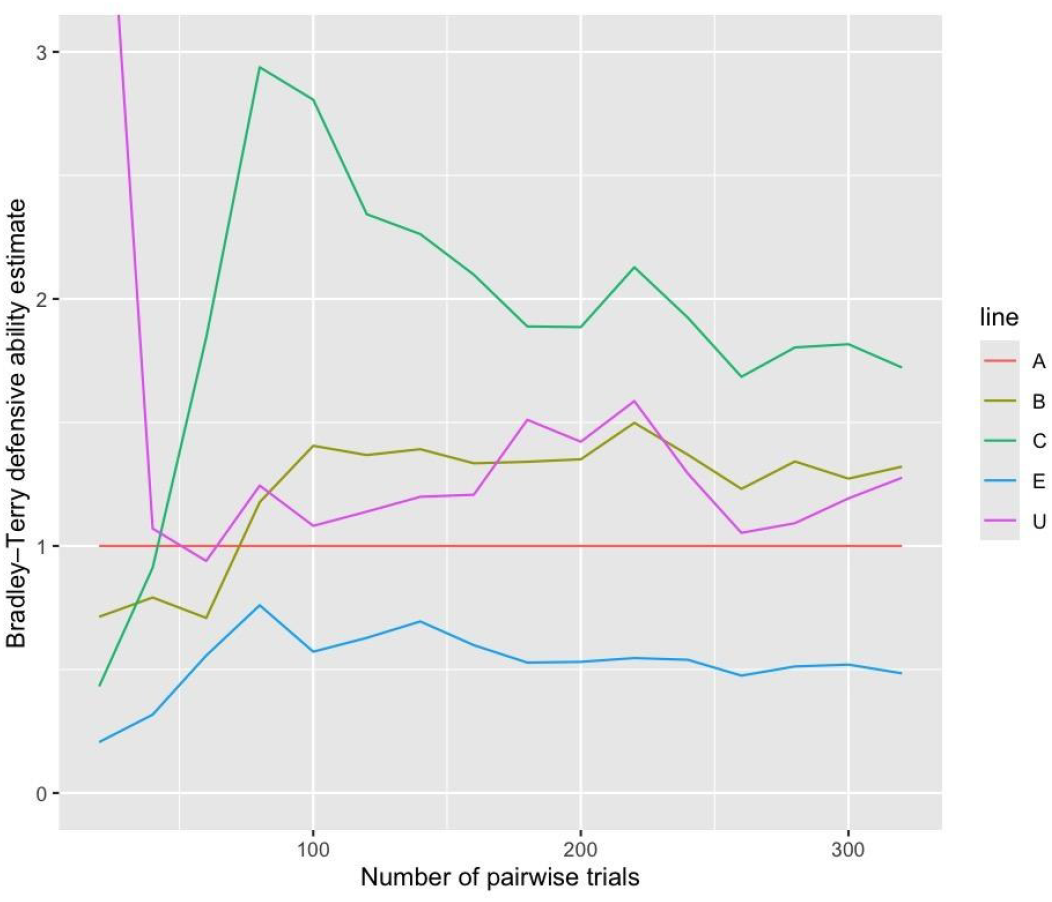
Sequential application of Bradley-Terry estimation following subsampling of the data. The figure shows how the estimates of offense (P2) abilities stabilize as the number of mating trials increases. The rank ordering of the five lines appears to stabilize with the last switch occurring after 225 mating trials.

## Discussion

When sperm competition assays use a single standard male tested against a series of focal lines, relative competitive ability is straightforward to quantify. In natural populations, however, males encounter many different competitors, creating a complex network of pairwise trials. It is not obvious *a priori* that the outcomes of these trials can be reduced to a simple ranking: for example, line X might outperform most lines except Y, while line Y may perform poorly against all lines except X. The Bradley-Terry model assumes that a transitive ranking exists and evaluates this assumption by estimating parameters that best predict observed outcomes of sperm competition.

In the data presented here, the Bradley-Terry model captures differences in sperm competitive ability and fits the data far better than a null model in which all lines are equivalent. Deviance statistics indicate an overall acceptable fit, although certain lines consistently show larger discrepancies between observed and predicted outcomes.

Figure 3 shows that line E deviates consistently the model’s predictions. An important application of this framework is to flag such outliers, including genetically modified lines that exhibit atypical competitive performance and may point to disruptions in processes such as seminal fluid protein function or sperm numbers.

A good fit to the Bradley-Terry model is consistent with Parker’s (1970) “fair raffle” hypothesis, in which paternity share is proportional to the number of sperm entering storage within the female. Under this model, males that contribute more sperm to storage have proportionally higher reproductive success, analogous to holding more raffle tickets. Consequently, the rank order of sperm competitive ability should reflect the rank order of sperm storage success among genotypes. The present experiment uses a small set of readily distinguishable lines (and their F1 progeny); extending this approach to a broader panel of wild-derived strains would provide a stronger test of the fair raffle model in a more natural context.

The model developed here does not incorporate cases in which sperm competition outcomes depend on female genotype. We have previously shown that the rank order of male sperm competitive success varies across female genotypes, indicating that females play an active and genetically variable role in mediating these outcomes.

Female genotypes differ in their responses to ejaculate components, influencing sperm storage, utilization, and ultimately paternity, consistent with variation in the “receipt” of signals associated with ejaculate quality (Clark and Begun 1998). Quantitative trait locus analyses have identified genetic factors underlying female refractoriness and sperm competition (Civetta et al. 2000; Chow et al. 2010). More broadly, sperm competition is an interaction phenotype: outcomes depend on specific combinations of male and female genotypes rather than intrinsic male ability alone (Clark, Begun, and Prout 1999). This view is reinforced by evidence that females modulate sperm dumping and sperm use following multiple mating (Snook and Hosken 2004), as well as by studies in *Drosophila pseudoobscura* demonstrating female-mediated sperm death and differential sperm survival (Holman and Snook 2008). Together, these processes highlight a coevolving system in which male × female interactions help maintain genetic variation and complicate predictions of evolutionary dynamics in natural populations.

The sensitivity of male rank order to female genotype underscores the mechanistic complexity of sperm competition. Despite substantial progress, the underlying mechanisms in *Drosophila* remain incompletely understood, although ongoing work continues to identify the genes and pathways involved. The Bradley-Terry framework developed here provides a useful step toward this goal by offering a quantitative approach for predicting sperm competition outcomes, while also highlighting where additional biological complexity must be incorporated. Moreover, the use of an order-dependent Bradley-Terry framework will have broad application to any pairwise competition that includes asymmetry between the contestants. For example, sperm competition assays could be scaled up to a large collection of genotypes. After double-matings are done, researchers would extract and bar-code the DNA from progeny, combine those into a single library, and determine paternity proportions for all the crosses in a single sequencing run. Two-order Bradley-Terry analysis would then determine the sperm competition “winners” and “losers” for all combinations, and they could be associated with attributes of the lines. Genetic lines that are outliers in the analysis could also be informative about factors driving reproductive success.

## Acknowledgements

We thank Ben McCormick and Dan Barbash for sharing the A and B Drosophila lines and members of the Clark and Wolfner labs for discussion. This work was supported by NIH grant R01 HD059060 to AGC and MFW.

## Supplemental Figures

**Figure S1:**
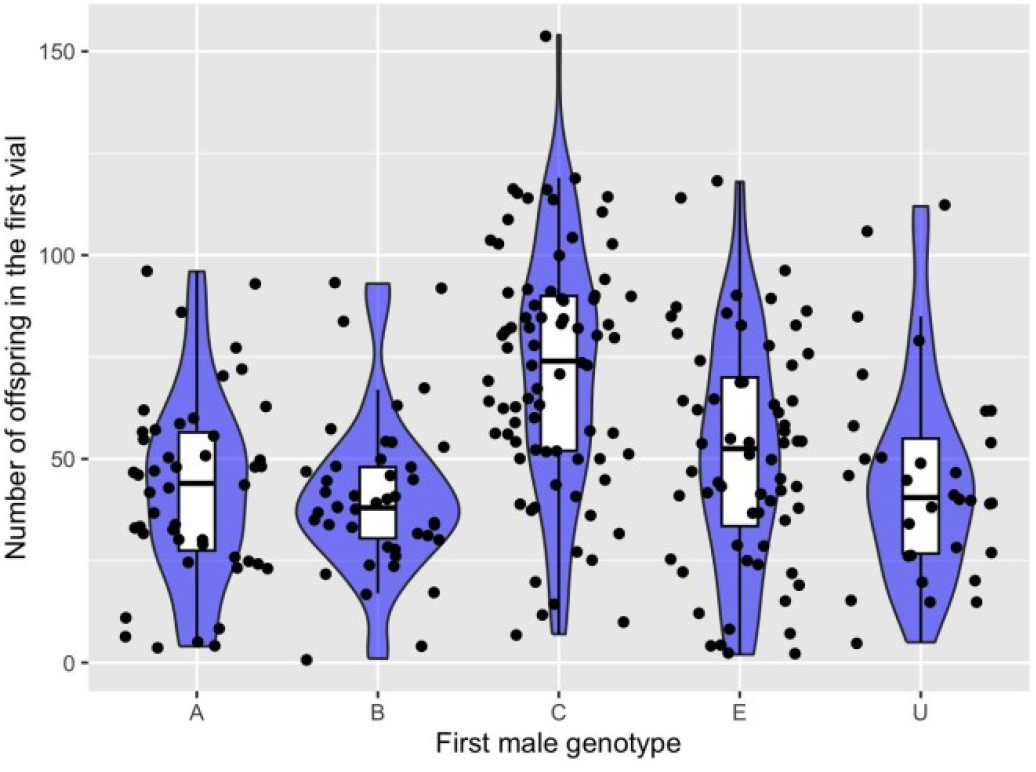
Number of offspring sired by each genotype in the first vial 0

**Figure S2:**
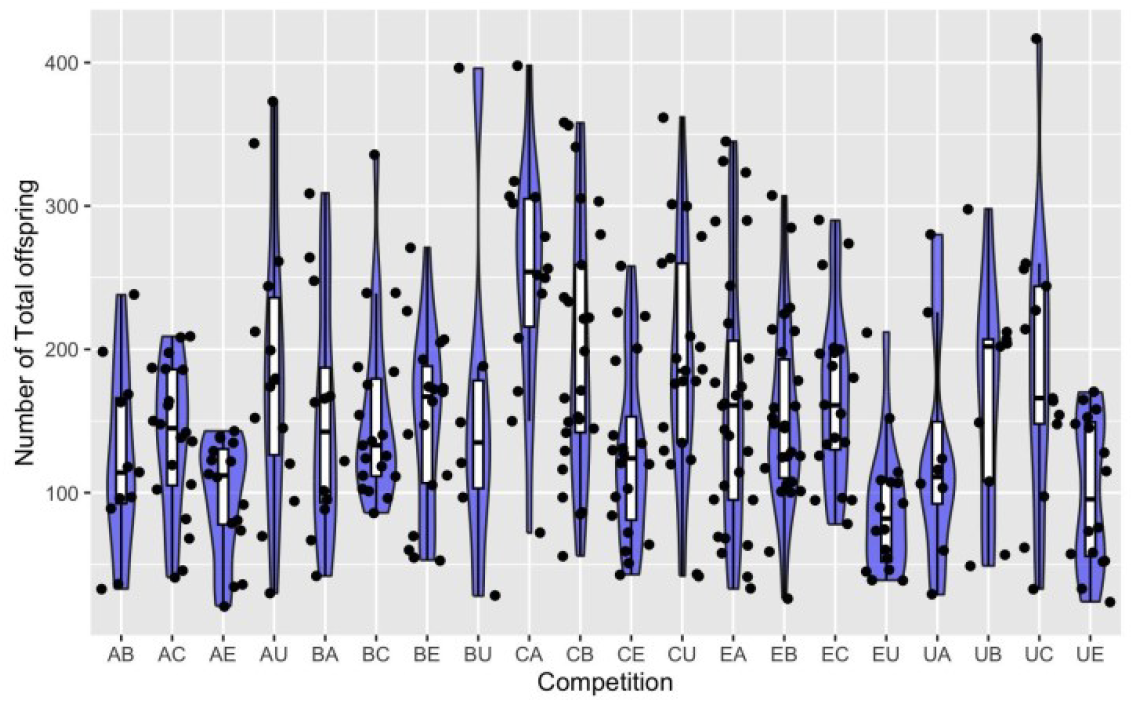
The number of offspring sired within each of the 20 distinct double-mating trials.

**Figure S3:**
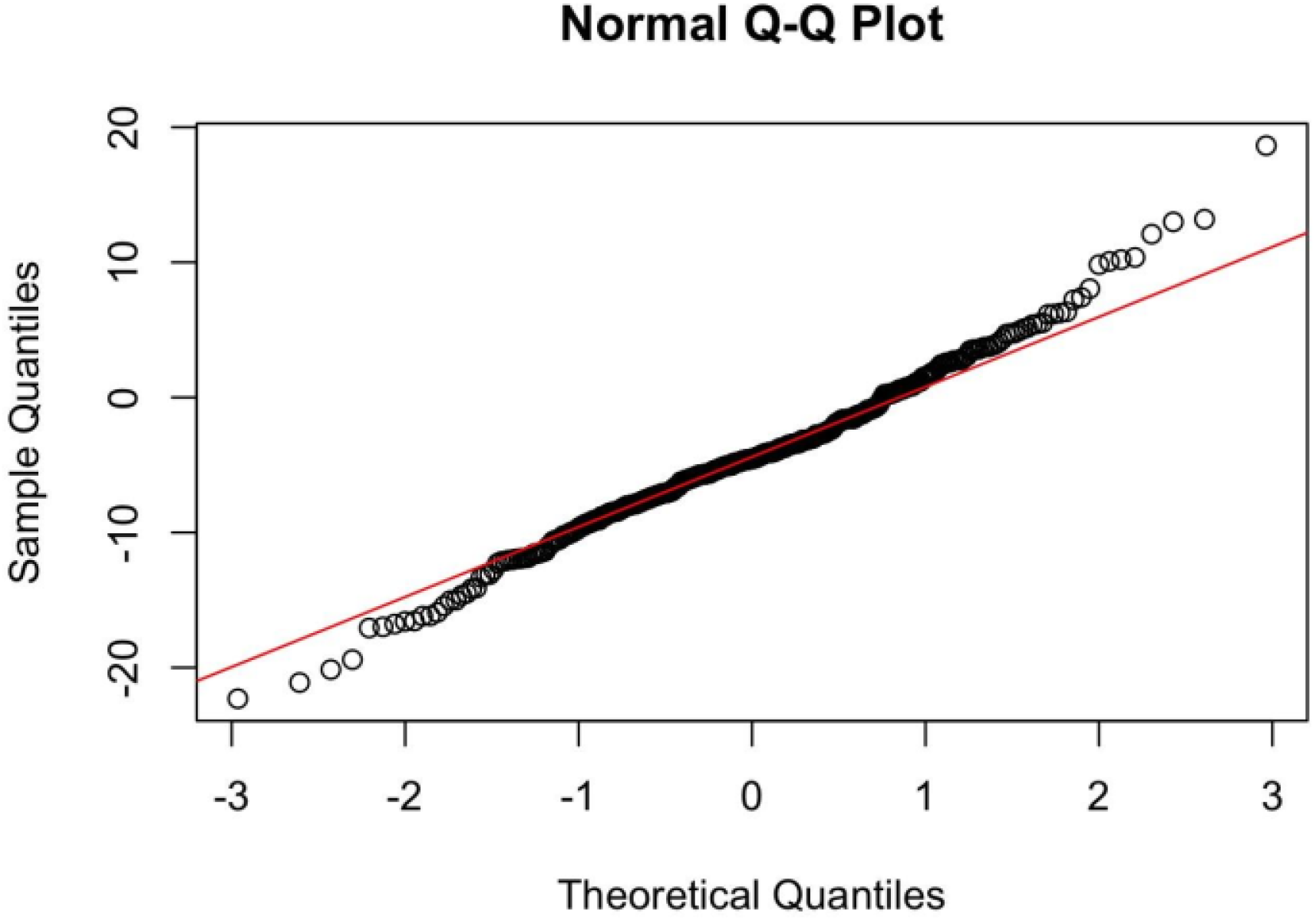
The Q-Q plot for the basic Bradley-Terry model.

